# Cryo-EM structure of the mycobacterial 70S ribosome in complex with ribosome hibernation promotion factor RafH, reveals the unique mode of mycobacterial ribosome hibernation

**DOI:** 10.1101/2023.04.18.537051

**Authors:** Niraj Kumara, Shivani Sharmaa, Prem S. Kaushala

## Abstract

Ribosome hibernation is a key survival strategy bacteria adopt under environmental stress, where a protein, hibernation promotion factor (HPF), transitorily inactivates the ribosome and slows down its overall protein synthesis. The mechanism is well studied in enteric bacteria, which mainly hibernate its ribosome in 100S disome form through a dual domain, long HPF (HPF^long^) or a single domain, short HPF (HPF^short^) in concert with another ribosome modulation factor. Mycobacteria under hypoxia (low oxygen) stress overexpresses RafH protein regulated under DosR regulon, a critical factor for its survival. The RafH, a dual domain HPF, an orthologue of bacterial HPF^long^, hibernates ribosome in 70S monosome form only. Here we report the cryo-EM structure of *Mycobacterium smegmatis*, a close homologue of *M. tuberculosis*, 70S ribosome in complex with the RafH factor at an overall 2.8 Å resolution. The RafH N-terminus domain (NTD) is conserved and binds to the decoding center of the ribosomal small subunit, a similar binding site of HPF^long^ NTD, but additionally it also interacts with the inter subunit bridge, B2a. Contrary to the HPF^long^ CTD, the RafH CTD, which is larger, binds to a unique site at the platform binding center of the ribosomal small subunit and sandwiches between bS1 and uS11 ribosomal proteins. The two domain connecting linker regions, which remain mostly disordered in earlier reported HPF^long^ structures, interacts mainly with the anti-Shine Dalgarno sequence of the 16S rRNA. The helix H54a of 23S rRNA, unique to the mycobacterial ribosome, adopts a different conformation and come close to RafH CTD, suggesting its role in ribosome hibernation. RafH inhibits *in-vitro* protein synthesis in a concentration dependent manner. Further, the modeling studies provided the structural basis for the incompatibility of mycobacterial ribosomes forming 100S like hibernating ribosomes.

## Introduction

*Mycobacterium tuberculosis* (Mtb) is the causative agent of one of the most deadly bacterial disease, tuberculosis (TB), which remains the second leading cause of death from a single infectious agent after COVID-19. The COVID-19 pandemic has further worsened the decade’s effort on TB control and made it challenging to achieve WHO’s goal of “ending the global TB epidemic” by 2030 (WHO’s Global Tuberculosis Report 2022)^1^. The situation remains worrying with the emergence of multi-drug resistance (MDR) and extensive drug-resistant (XDR) TB strains, resistant to the currently used drugs to treat TB infection^2^.

The Mtb’s long persistence is attributed in part to a minor subpopulation of non replicating, drug-tolerant persisters cells, capable of surviving in the hostile environment of the host macrophages in a dormant state without eliciting a host immune response, known as latent Tuberculosis infection (LTBI)^3–6^. An estimated one-quarter of the world’s population possesses dormant Mtb, and 5-10 % develop acute tuberculosis infection^7–9^. Therefore, the dormant Mtb serves as a vast reservoir for TB infection^10^. During latent infection, protein synthesis or translation, a vital cellular process that consumes nearly half of the cell’s energy^11^ and 40% of known antibiotics inhibit specific translation steps^12^, is globally downregulated. Therefore, Mtb becomes less susceptible to ribosome targeting antibiotics^13^.

Ribosome hibernation, in which a protein factor inactivates the ribosome by reversible binding, is a highly conserved, tightly regulated process in bacteria and responsible for the survival of growth-arrested bacterial cells under environmental stresses in a drug-tolerant state^14–16^. The hibernating ribosomes appear to be less susceptible to degradation by RNase^17^.

The process is well studied in enteric bacteria, which possess two forms of HPF. The HPF short (HPF^short^), a single domain protein, induces 100S ribosome (disome) formation with another factor known as ribosome modulation factor (RMF)^18, 19^. The HPF long (HPF^long^), a two domain protein factor, is solely responsible for inducing the 100S ribosome formation. The HPF^long^ induces 100S disome formation through dimerization of its C-terminus domain^20, 21^. The molecular mechanism of 100S ribosome hibernation is thoroughly studied^15^, and the cryo-EM structures of hibernating 100S ribosomes from different bacterial species are available^18–24^. Another mode of ribosome hibernation is induced by a single domain protein, the protein Y (encoded by gene *yfiA*, also known as *raiA*)^25^ and its orthologue in chloroplast ribosome, PSRP-1^26, 27^ which hibernates ribosome in the 70S (monosome) form only.

Although the overall translation machinery in mycobacteria is conserved, however, there are unique structural features associated with the ribosome architecture^28–36^ such as H54a, ∼110 nucleotide insertion in H54 of the 23S rRNA^28–30^. Another distinctive feature associated is its ribosome hibernation which has been proposed to be a primary survival mechanism for non replicating Mtb ^16, 31^. Mycobacteria hibernates ribosomes in 70S monosome form only, any higher order ribosome structure, such as 100S disome, are not reported so far^31, 32, 37^.

Mycobacterial HPF, the mycobacterial protein Y (MPY), (also designated as a ribosome associated factor under stasis RafS) induces 70S ribosome hibernation under different environmental stress, such as low carbon^37^, low Zn^++^ ion^31^, and during stationary phase^32^. The MPY possesses two domains and a connecting linker region^16^, a typical dual domain, HPF^long^ like organization^15^. The MPY CTD and linker region remains disordered in reported structures^31, 32^. Thus, its binding site information and the structural basis of MPY’s inability to induce ribosome dimerization to form 100S remains unknown.

Mycobacterium contains another HPF known as RafH, which is overexpressed under hypoxia stress and regulated by the DosR regulon. It’s believed that Mtb encounters multiple stresses, primarily low oxygen content, the hypoxia, in host macrophages. Hypoxia induces the DosR regulon, which overexpresses nearly 48 genes, including, RafH (ribosome associated factor under hypoxia)^37^ expressing gene MSMEG_3935 in *M. smegmatis* and Rv0079 in *M. tuberculosis*^38^. The RafH is a hibernation promotion factor (HPF) that induces ribosome hibernation and stabilizes ribosomes in the 70S (monosome) form^37^. It promotes cellular viability in a growth-arrested state and appears to be the major factor responsible for Mtb’s survival under hypoxia^37^. Overexpression of the factor led to an early entry to the stationary phase in *E. coli,* and its gene was found to be conserved in many clinical isolates^39^. RafH, an orthologue of the HPF^long^, also possesses two domain architecture, but still cannot induce ribosome dimerization and 100S formation as reported for other similar HPFs. However, the structural basis of RafH induced ribosome hibernation and its inability to form 100S-like disome is unknown, as no structure is available.

Here, we report the single particle cryo-EM structure of the *M. smegmatis* (a close homolog of *M. tuberculosis*) 70S ribosome in complex with RafH at an overall 2.8 Å resolution. In addition, we also report 70S ribosome in complex with RafH and bS1 ribosomal (r-) protein and 70S ribosome in complex with RafH and E-site tRNA, both cryo-EM structures at 3.5 Å resolution. The structure reveals that RafH NTD binds to the conserved binding cleft between the head and body at the decoding center of the ribosomal small subunit. Its binding overlaps with the binding sites of translation factors. It also interacts with the inter subunit bridge, B2a. Whereas RafH CTD, which is larger, and has a repeated HPF^long^ CTD like domain architecture, binds to a unique position at the platform binding center of the small subunit, which has not been reported before. The linker region connecting two domains interacts primarily with the anti-Shine Dalgarno (a-SD) sequence of the 16S rRNA. Intriguingly, the study reports the remarkable interaction between the HPF linker and a-SD in atomic details for the first time. The *in-vitro* translation assay showed, RafH inhibits protein synthesis in a concentration dependent manner. Further, modeling studies provided the structural interpretation for RafH’s inability to induce ribosome dimerization and formation of 100S-like ribosome structures, known to be induced by the HPF^long^.

## Results

### 70S ribosome RafH complex formation and protein synthesis inhibition

The 70S ribosomes were purified by sucrose density gradient ultracentrifugation (Fig. 1a). To remove the co-purified P-site bound tRNA, the 70S ribosomes were dissociated into their respective subunits by lowering the MgCl_2_ to 1 mM (Fig. 1b) and further re-associated by incubating equimolar concentrations of 50S and 30S subunits in 20 mM MgCl_2_ (Fig. 1c). The 70S ribosome RafH complex, prepared by mixing the re-associated 70S ribosome with purified RafH protein, was confirmed by sucrose pelleting assay (Fig. 1d). The RafH protein band was visible in SDS-PAGE for the pellet fraction of the ribosome RafH reaction mixture. As expected, the corresponding band was absent in the pellet fraction of ribosome without RafH (Fig. 1d). The *in-vitro* translation assay, performed by titrating ribosomes with different stoichiometry ratios of RafH, showed that RafH inhibits the protein synthesis in a concentration dependent manner (Fig. 1e), which also confirmed that the purified protein was in an active conformation. Further, the cryo-EM grid preparation conditions were optimized. The cryo-EM image showed an even distribution of intact ribosome particles with optimum ice thickness (Fig. 1f).

**Figure 1.**
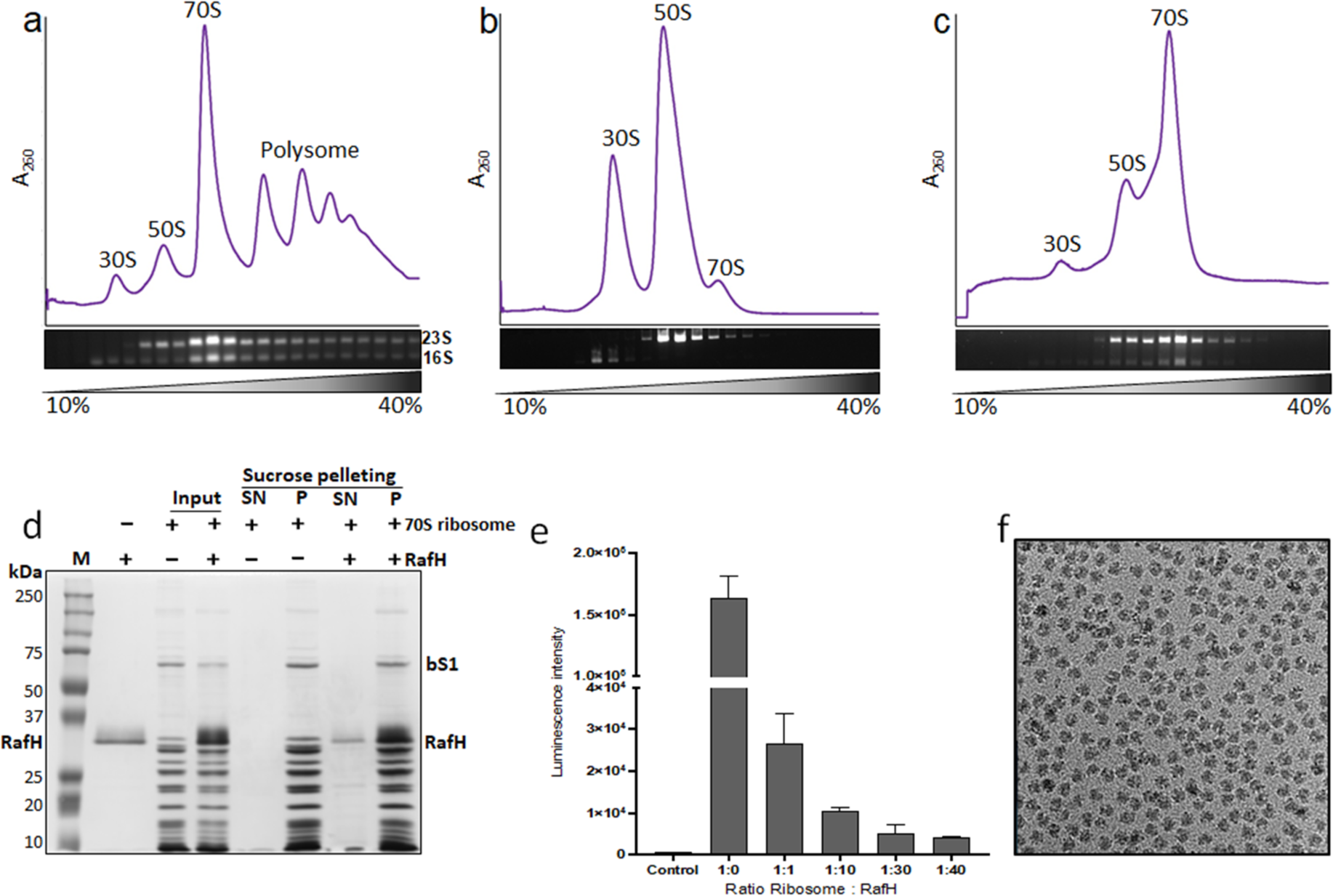
70S ribosome RafH complex. (a-c) The 10% to 40% sucrose density gradient fractionation profile and corresponding peaks analysis and agarose gel are shown for initial ribosome purification (a), after dissociation (b), and after re-association (c). (d) the 70 ribosomes RafH complex formation and sucrose density pelleting analyzed on 12% SDS-PAGE, lane 1 - marker, lane 2 - pure RafH protein, lane 3, 4 - input, lane 5 to 8 - supernatant and pellet fraction after pelleting on a sucrose cushion. (e) *In-vitro* protein synthesis assay by titrating ribosome and RafH at different stoichiometric ratios. (f) The 2D cryo-EM micrograph collected during the initial grid screening stage in JEOL 2200 FS microscope with a Gatan K2 Summit camera.

### Single particle reconstruction and sorting structural heterogeneity

For elucidating the molecular mechanism of mycobacterial ribosome hibernation, the structure of the 70S ribosome RafH complex was determined by single particle cryo-EM reconstruction using Relion-3.1.4. After extensive 3D classification, 1,53,262 particles were selected from the classes showing bound RafH. The 3D refinement yielded an initial cryo-EM map of 3.0 Å resolution. Further, the map quality and resolution was improved to 2.8 Å by doing CTF refinement and particle polishing (Supplementary Fig. S1a). To further improve the density for RafH CTD, these particles were subjected to signal subtraction with 3D classification without alignment in 5 classes. Class 1, showed fragment cryo-EM mass for RafH CTD, class 2, showed the presence of bS1 protein in addition to RafH CTD, class 3 showed only RafH CTD, class 4, showed the presence of E-tRNA with RafH CTD and the remaining 1% in class 5 were unaligned. The three classes 2, 3, and 4 all having cryo-EM density for RafH CTD, with a total 1,10,934 particles, yielded a 2.8 Å cryo-EM map after 3D refinement and postprocessing (Supplementary Fig. S1b). Multi-body refinement further improved the map quality and resolution to 2.7 Å and 2.9 Å for the LSU and SSU, respectively. A similar approach of multi-body refinement was applied to improve the quality of cryo-EM maps for classes 2 and 4 separately. These 3 maps, of ribosome RafH complexes (map1), with bS1 (map2), and with E-site tRNA (map3) were selected for model building and structure interpretation.

### Cryo-EM structure of 70S ribosome RafH complex

Overall, the cryo-EM map shows high resolution features (Fig. 2a, b) with distinctly visible secondary structures α-helices and β-sheets for RafH NTD (Fig. 2c). Most of the amino acid residue side chains and nucleotides were clearly visible in our cryo-EM map (Fig. 2c, d and Supplementary Movie 1, 2). The local resolution calculated using ResMap showed the resolution ranges from 2.5 Å to 5.5 Å, with most of the region having better than 3.5 Å resolution (Supplementary Fig. S2). Some of the flexible regions, such as RafH CTD, E-site tRNA, L1 stalk, L7/L12 stalk, bS1, uS2, and H54a, having a lower resolution, were interpreted by applying a low pass filter of 3.5 Å resolution to the final map (Fig. 2a, b and Supplementary Fig. S3). For bS1 r-protein, the two N-terminus domains, OB1 and OB2, were clearly visible, whereas other parts were disordered (Fig. 2b and Supplementary Fig. S3a, b).

**Figure 2.**
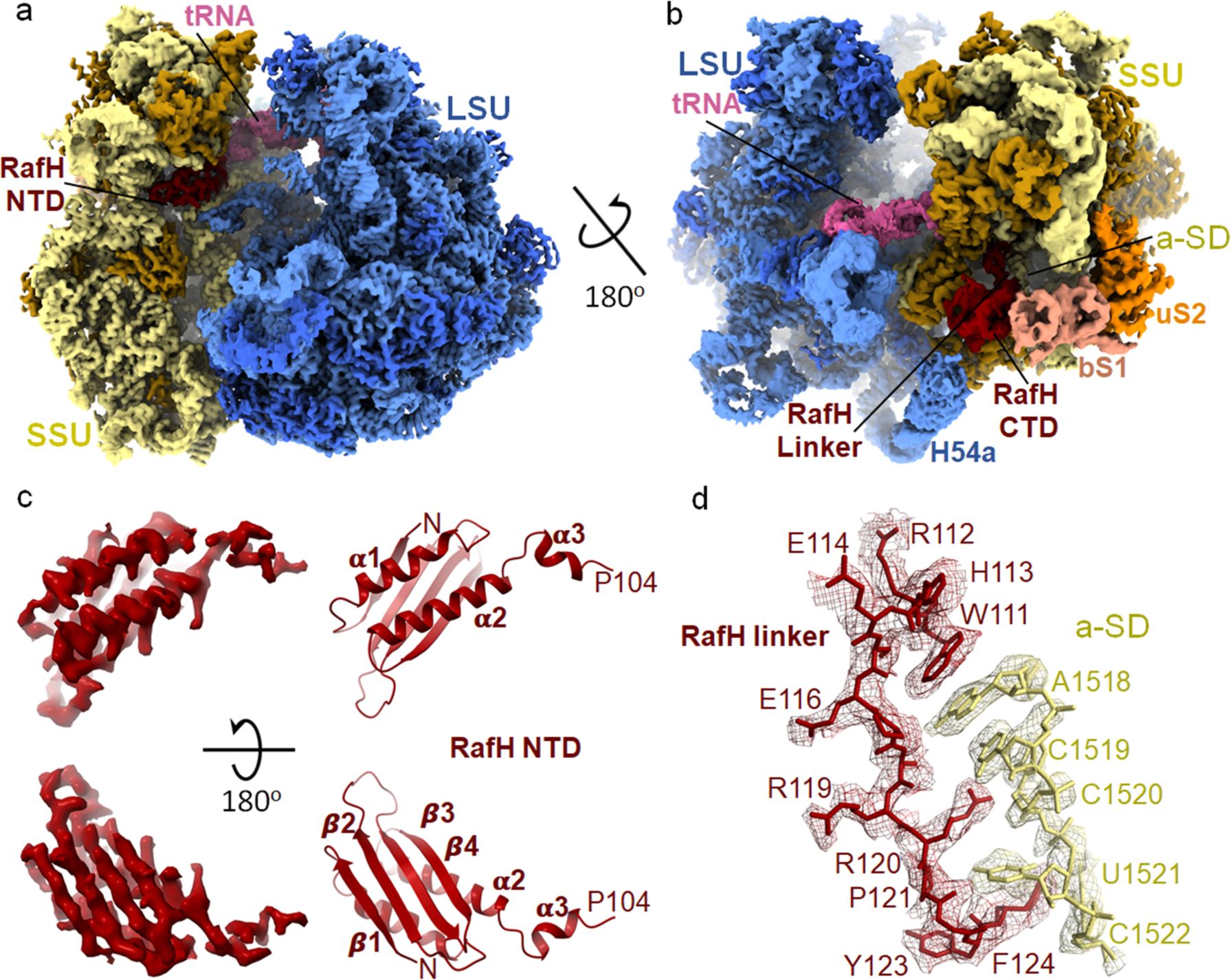
Cryo-EM structure of *Mycobacterium smegmatis* 70S ribosome RafH complex. (a, b) the overall architecture of the 70S RafH complex is shown in the mRNA entry site (a) and mRNA exit site (b) by a rotation through a diagonal axis. The SSU 16S rRNA (khaki), SSU r-proteins (dark golden), RafH (maroon), tRNA (pink), the LSU 23S rRNA and 5S rRNA (cornflower blue), LSU r-proteins (royal blue), bS1 (dark salmon) and uS2 (orange) are labeled. The single particle reconstruction data processing summary is shown in Supplementary Fig. S1, gold standard FSC and local resolution of final maps is shown in Supplementary Fig. S2, The cryo-EM maps for individual r-proteins, bS1 and uS2 and E-site tRNA and their model is shown in Supplementary Fig. S3, a full RafH model is shown in Supplementary Fig. S4. (c) the RafH NTD cryo-EM density (left panel) and model in ribbon (right panel), the top panel is rotated by 180°C along X-axis, and shown in the bottom panel, the secondary structures are labeled. (d) the cryo-EM density in mesh and model in stick style corresponding to RafH linker region residues, 111-124 (maroon) and a-SD region of 16S rRNA nucleotides, 1518-1522 (khaki), are shown. For more clarity, an animation is provided Supplementary Movie 1.

### RafH NTD binds to the conserved binding pocket in the 70S ribosome

RafH is a ∼30 kDa protein with 258 amino acid residues. It possesses two domains, the N-terminus domain (NTD), residues 1-100, C-terminus domain (CTD), residues 131-258, and these two domains are connected by a flexible linker region, residues 101-130 (Supplementary Fig. S4). The RafH NTD has a conserved domain having α/ý fold with ý_1_α_1_ý_2_ý_3_ý_4_α_2_ topologies where 4 β-strands form an antiparallel β-sheet and the two α-helices stacks to the one side of the β-sheet. A mini helix, α_3,_ connects the RafH NTD through a small loop (Fig. 2c and Supplementary Fig. S4). RafH NTD binds to the cleft between the head and body of the SSU (Fig. 2a and Fig. 3) to a similar binding site reported for HPF^long^ NTD, HPF^short,^ and YfiA in ribosome structures^15^. At this cleft, the NTD makes extensive interactions with the 16S rRNA, r-proteins uS9, and also with the inter subunit bridge B2a (Fig. 2a, Fig. 3, Supplementary Fig. S5 and). Some predominant interactions are shown in Fig. 3. The side chain of residue R75 of the helix α2 interacts with A1477-G1478 of 16S rRNA and A2137 of 23S rRNA (Fig. 3 and Supplementary Movie 2), thus providing more stability to 70S as these nucleotides form an inter subunit bridge, B2a^40^. The positively charged side chain residues, K21, R24, and R28 of the helix α1 make electrostatic interaction with the backbone phosphate of the h44 of 16S rRNA, residues G1478, U1479, C1480, G1481. The A770 of h24 interacts with the H30 of RafH (Fig. 3). The arginine-rich patch of α2 composed of R84, R88, and R91 interact with the C1382, C1383, and G1384 of the 16S rRNA (Fig. 3). Similarly, the residues R37, R39 of ý2 strands, residue Q55 of ý3, and R66 of ý4 forms a positively charged patch that stacks against U947 and G948 of h31 of the 16S rRNA (Fig. 3). The W96, which harbors in the mini helix α3, makes stacking interaction with the G673 of h23 (Fig. 3). The RafH NTD also interacts with the r-protein uS9. The H35 and D59 residues of RafH interact with the C-terminus residue R150 of the uS9 r-protein, whereas the N64 residue of RafH interacts with the K149 of the uS9 r-protein.

**Figure 3.**
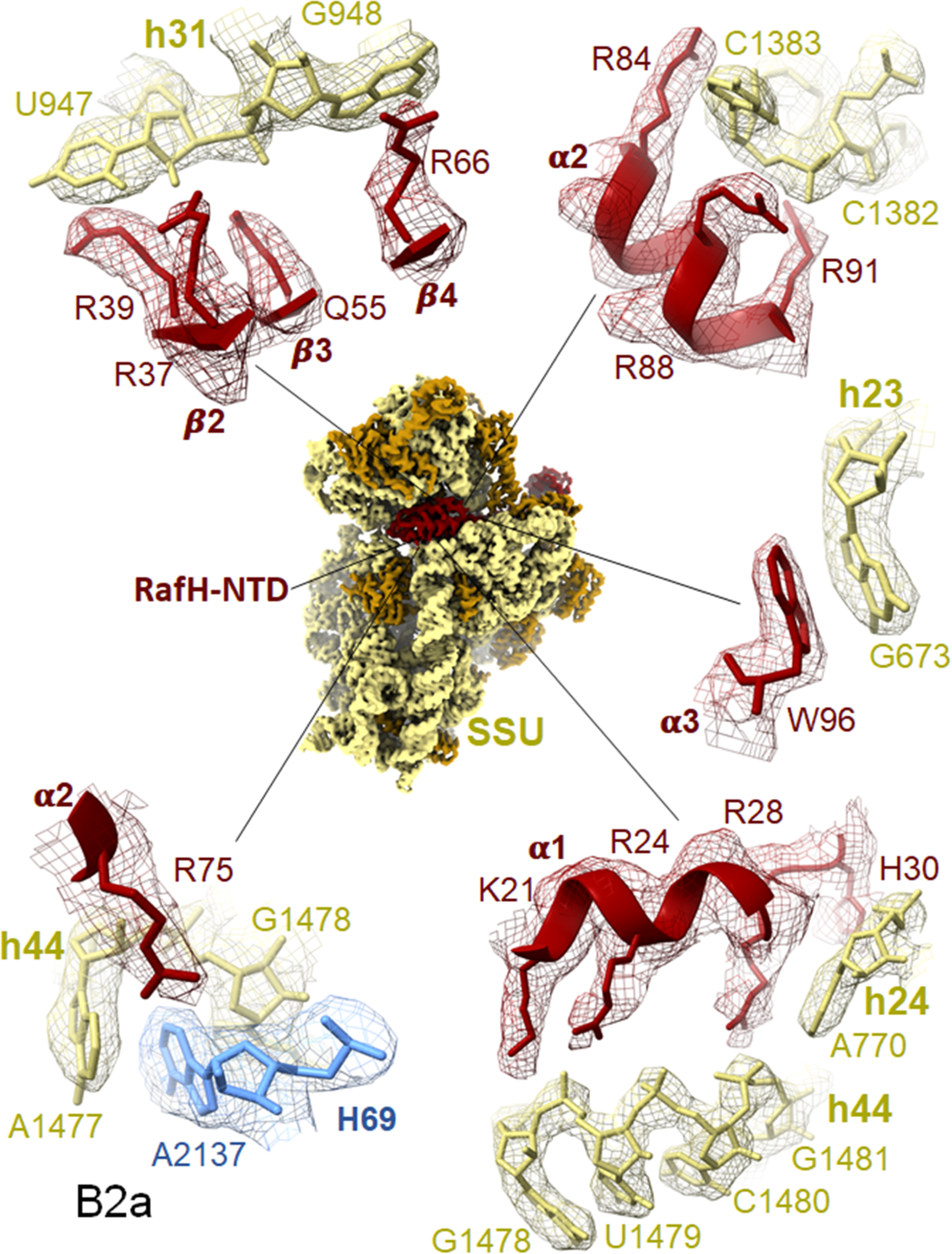
Ribosome and RafH NTD interaction. The cryo-EM density in surface view for the small subunit with RafH at the centre, and its magnified regions where the cryo-EM density in mesh and model in stick and ribbon are shown. For clarity, the ribosomal large subunit is not shown. The RafH 16S rRNA interaction in counterclockwise from the bottom left for α1 R75 with Bridge B2a, α1 with h44, α3 W96 with h23 G673, α2 with C1382-C1383, residues from β2, β3, and β4 with h31 U947, G948 are shown. For more detail see Supplementary Movie 2, See Supplementary Fig. S5, Supplementary Table S2, and text.

### RafH CTD binds to the unique binding site in the 70S ribosome

In the cryo-EM map, the resolution for the RafH CTD was relatively low compared to its NTD (Fig. 2a, b and Fig. 4a, b). However, the RafH CTD model obtained from AlphaFold2 nicely docked in the cryo-EM density designated to RafH CTD (Fig. 4b). We could clearly see α-helices and the side chains for some of the residues; R215, E219, R220, L221, and L223 for one of the a α-helices, α5 (Fig. 4b), which has further confirmed its binding site. The RafH CTD binds to the mRNA ‘platform binding center (PBC)’ composed of proteins bS1, uS7, uS11 and bS18 with 16S rRNA helices h26, h40 and 23S rRNA helix H54a. The uS11 and OB2 domains of the bS1 sandwich the RafH CTD (Fig. 2b, Fig. 4a). RafH CTD is composed of two similar protein folds, a α helix with 4 stranded antiparallel ϕ3-sheet, having ϕ3_5_α_4_ϕ3_6_ϕ3_7_ϕ3_8_α_5_ϕ3_9_ϕ3_10_ϕ3_11_ϕ3_12_ topologies. The ϕ3_6_ϕ3_7_ϕ3_8_ϕ3_9_ forms a 4 stranded antiparallel ϕ3-sheet where α4 stacks to one side of the ϕ3-sheet and form the first protein fold. Similarly, ϕ3_5_ϕ3_10_ϕ3_11_ϕ3_12_ forms another 4 stranded antiparallel ϕ3-sheet and α5 stacks to one side of it to form the second protein fold. The two folds are connected by a loop region (Supplementary Fig. S4). The ϕ3-sheets of each fold stack nearly parallel to each other and form a dimer-like structure^41^ (Fig. 4b and Supplementary Fig. S4). On the contrary, the HPF^long^ CTD (enteric bacteria) is composed of a single protein fold (Supplementary Fig. S6) but attains a RafH CTD like architecture by its dimerization (Fig. 4c), as a consequence of which 100S disome^15^ formation takes place (Fig. 4b, c).

**Figure 4.**
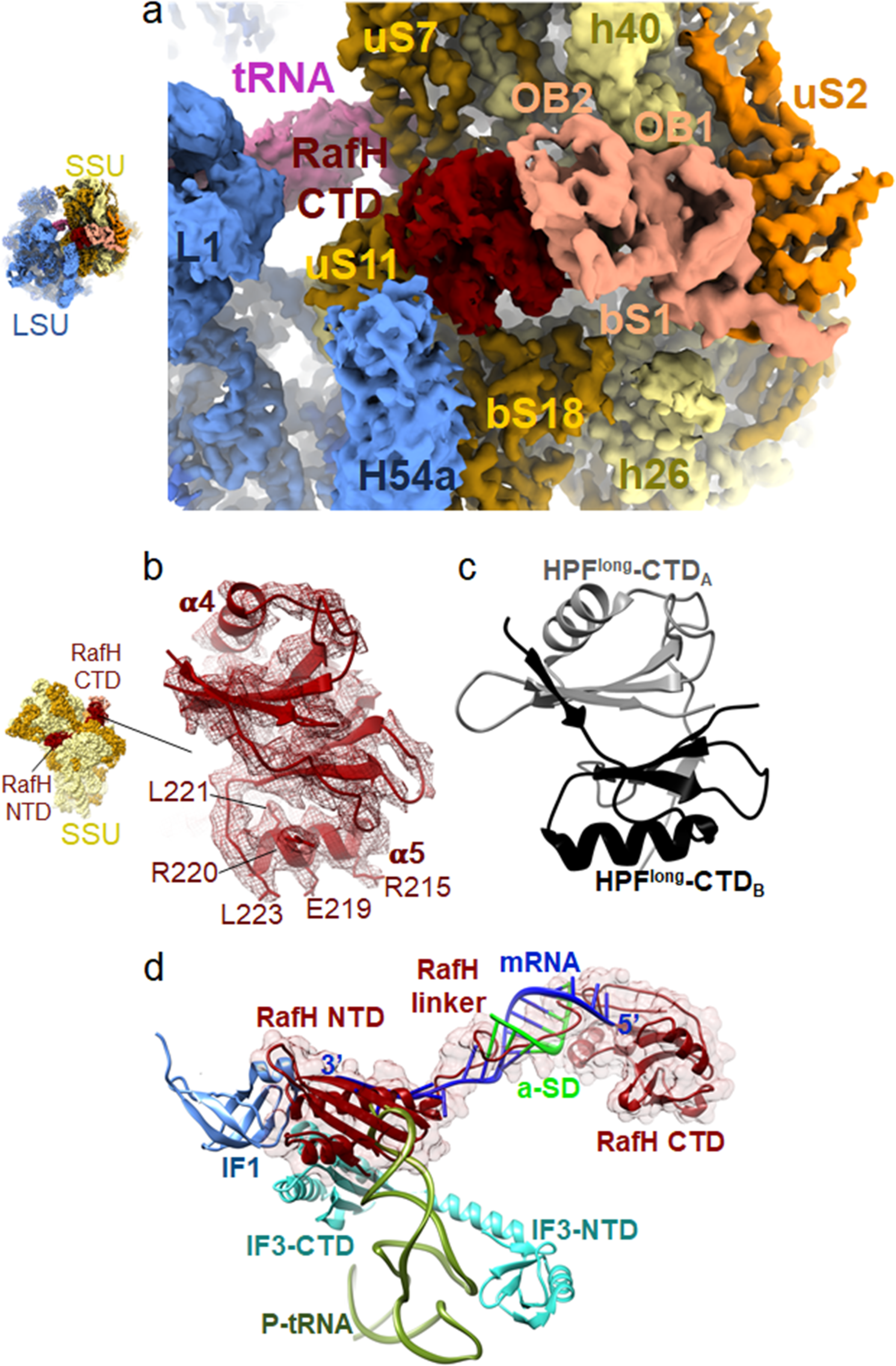
RafH CTD structure and its binding site on the ribosome. (a) The RafH CTD binding site environment in cryo-EM map in the surface style for 70S RafH complex is shown, and the same color scheme of Fig. 2a, b is used. A thumbnail for the 70S is shown on the left. (b) cryo-EM density corresponding to RafH CTD in mesh, model in ribbon, and stick is shown. The thumbnail is shown on the left. (c) the structure of HPF^long^ CTD dimer (PDB ID; 6T7O) with its first monomer (A) (gray) and second monomer (B) (black) are shown. (d) the pre-translation initiation structure SSU (PDB ID; 5LMT) docked onto the ribosome RafH complex SSU structure. For clarity, only the RafH in ribbon and 95% transparent surface, initiation complex factors: mRNA (blue), a-SD (green), IF1 (cornflower blue), IF3 (cyan), and P-tRNA (dark olive green) are shown.

### The RafH linker interacts with the anti-Shine Dalgarno region of 16S rRNA

A flexible linker connects the RafH NTD and CTD with residues, stretch from 101 to 130 (Supplementary Fig. S4). The linker residues between W111 to F124 extensively interact extensively with the nucleotide stretch, A1518 to C1522, which harbors the anti-Shine Dalgarno region of the 16S rRNA (Fig. 2b, d and Supplementary Movie 1). This remarkable interaction involves residue W111 making a stacking interaction with the A1518 of 16S rRNA. The R120 side chain making electrostatic interaction with the C1519 base and phosphate of the C1520. The main chain of A118 also interacting with the nitrogenous base of A1518. The main chain of P121 interacting with U1521. The F124 making stacking interaction with C1522 of 16S rRNA (Fig. 2d). Altogether, these interactions would block the mRNA SD sequence interaction with the a-SD sequence of 16S rRNA, thus resulting in the inhibition of translation initiation. The linker also interacts with the anticodon stem-loop of the tRNA bound to the E-site of the ribosome (Fig. 2a, b). However, in the cryo-EM map, we could not see resolved nucleotide, maybe the cryo-EM density for E-site tRNA is from averaged tRNA, as the E-site tRNA co-purified during ribosome purification.

### RafH CTD and H54a of 23S rRNA would prevent the 100S ribosome formation in Mycobacteria

To understand the structural basis of RafH’s inability to induce ribosome dimerization resulting in 100S formation, molecular modeling, and docking were performed. The atomic coordinate of the 70S ribosome RafH complex was docked on each 70S monomer of the *Staphylococcus aureus* 100S ribosome dimer (PDB ID; 6FXC). It was found that the RafH CTD binds to a unique position near the uS11 r-protein and is surrounded by OB2 of bS1 and H54a of 23S rRNA in mycobacterial 70S ribosome. Besides this, OB1 of bS1 interacts with the uS2 r-protein (Fig. 5a). But, HPF^long^ CTD binds in the same vicinity too, close to the uS2 r-protein (Fig. 5b). Therefore, its binding site overlaps with the binding site of the bS1 r-protein, particularly its OB1 domain in mycobacterial 70S hibernating ribosome (Fig. 5a). As consequences of this, *M. smegmatis* 70S (monomer) in an *S. aureus* 100S like dimer architecture would have severe steric clashes at the dimer interface (Fig. 5c) where RafH CTD, H54a and bS1 OB1 of one ribosome would make steric clashes with h40, uS2, and bS1 OB2 of the second ribosome, and vice versa, (Fig. 5c). Therefore, would hinder the formation of a dimer of the ribosome. In contrast, the 100S formation is mainly stabilized by HPF^long^ CTD dimerization. In addition, uS2, bS18, h26, and h40 also stabilize the 100S dimer interface in some species (Fig. 5d)^15^.

**Figure 5.**
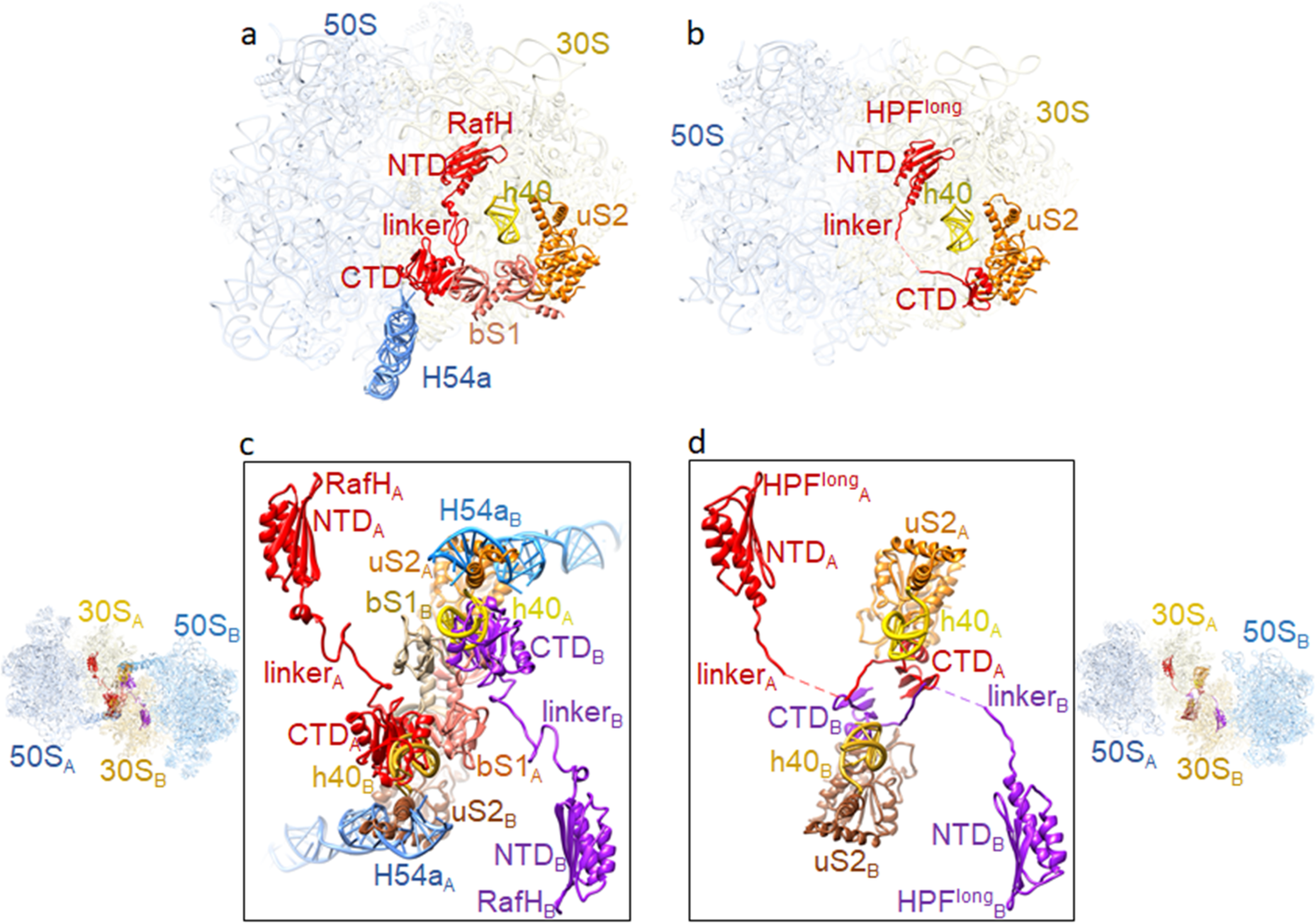
Comparison of RafH binding in 70S ribosome with HPF^long^ binding in 100S ribosome. (a) RafH, bS1, uS2, and h40 of 16S rRNA and H54a of 23S rRNA are shown with LSU and SSU in the 95% transparent background. (b) the corresponding position of HPF^long^, uS2, and h40 of 16S rRNA in one of the ribosomes of the *Staphylococcus aureus* 100S structure (PDB ID; 5NGM) is shown with LSU and SSU in the 95% transparent background. (c) two 70S ribosome RafH complex structures docked into the corresponding positions in *Staphylococcus aureus* 100S dimer structure (PDB ID; 6FXC) and RafH CTD interacting components are shown in 80% transparent background on the left side and magnified view with a white background are shown in the box on the right side. One 70S ribosome is labeled as A, and the other 70S ribosome is labeled as B. (d) the HPF^long^ interacting components uS2 and h40 of 16S rRNA in *Staphylococcus aureus* 100S ribosome dimer interface (PDB ID; 6FXC) are shown on the right side with 30S and 50S with 80% transparent background, and a magnified view with white background is shown in the box on the left side. Similar to (c), one 70S ribosome is labeled as A, and the other 70S ribosome is labeled as B.

## Discussion

This structural study reveals a unique mode of mycobacterial ribosome hibernation by RafH, a hypoxia induced HPF. The physiological significance of the RafH has been reported earlier by Trauner *et al.*, 2012^37^. RafH being a dual domain HPF, forms hibernating 70S ribosomes. The RafH NTD is conserved and binds to the decoding center of the ribosomal small subunit. Whereas RafH CTD is comparatively larger, already having a repeated HPF^long^ CTD like topology, and forms a similar dimer-like architecture as reported for the HPF^long^ CTD, which is required for 100S ribosome formation. Therefore, further RafH CTD dimerization is not possible; hence, RafH forms a hibernating 70S monosome only. Additionally, the H54a and bS1 would also prevent the mycobacterial ribosomes from forming a 100S like architecture. The RafH binding would block all known critical sites of translation initiation: the decoding center, the a-SD sequence of 16S rRNA, and the bS1 protein at the platform binding center of the ribosomal small subunit therefore, it inhibits protein synthesis. Thus, RafH has a distinct mode of ribosome hibernation (Fig. 6).

**Figure 6.**
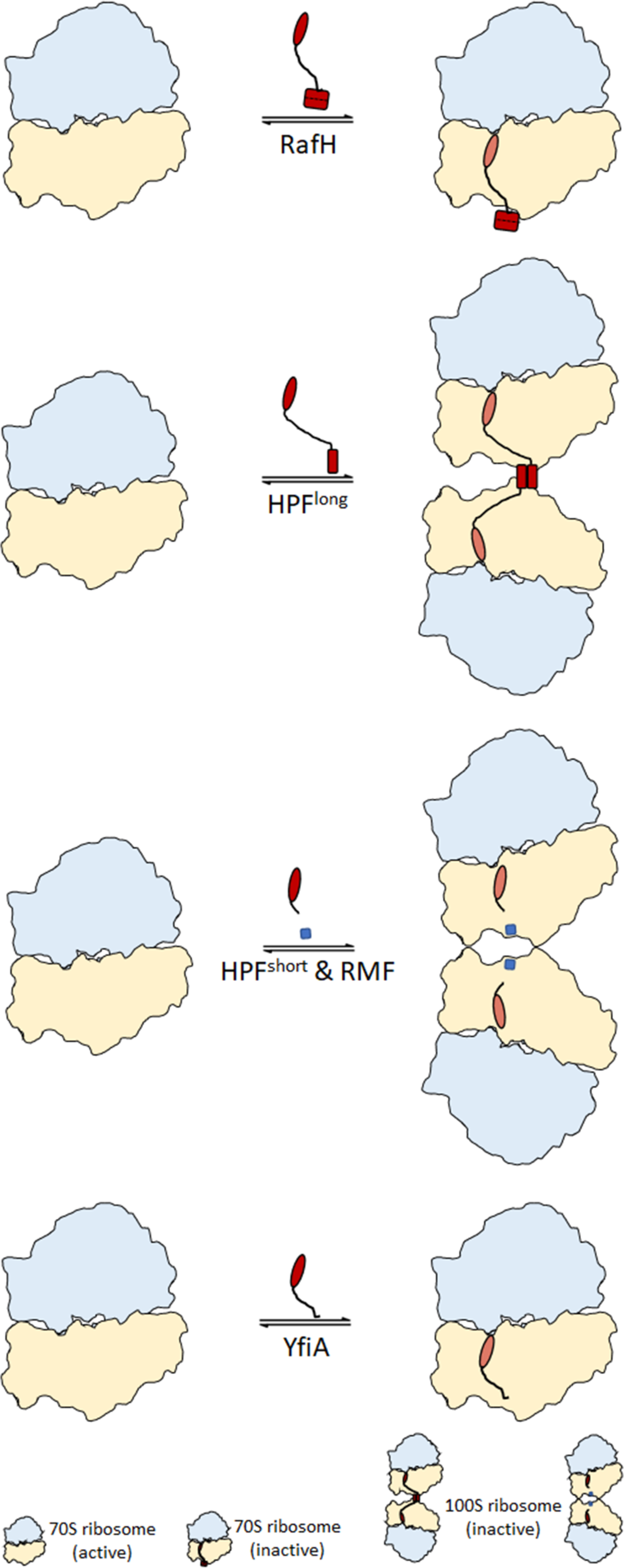
Different modes of ribosome hibernation. A schematic presentation for the different modes of ribosome hibernation. Top, RafH mediated hibernation in 70S form (from this study). Second from top, HPF^long^ induces ribosome dimerization and formation of 100S disome^20–24^. Third, from the top, HPF^short^ and RMF^18, 19^ induce ribosome dimerization and 100S ribosome formation. Bottom, YfiA hibernates ribosome in the 70S form^26, 27^.

The RafH NTD possesses a conserved structural fold and binds to a similar binding site to that of HPF^long^ NTD, HPF^short^, and YfiA^14, 15^. RafH NTD interacts with SSU predominantly through electrostatic interactions. But, the additional interaction of RafH R75 residue with the inter subunit bridge B2a (Fig. 3 and Supplementary Movie 2) suggests that the RafH binding further stabilizes the inter subunit interaction. The bridge B2a is also known to involve in the translation initiation and translational processivity in addition to strengthening the inter subunit interaction^42^. The RafH NTD binding would impede the binding of translation initiation factors IF1 and IF3, mRNA, and tRNAs at A and P sites (Fig. 4d). Besides this, the RafH NTD binding would also explicitly blocks the recruitment of leaderless (without 5’ UTR) mRNA^43^ onto the ribosomal small subunit and consequently blocking its translation initiation too. It is a notable observation because in Mtb, nearly 25% of mRNAs are the leaderless mRNA^44^, and it is believed that Mtb switches to the leaderless mRNA translation over leadered (with 5’ UTR) mRNA, as a survival strategy under stress^45^. The ribosome targeting antibiotics, particularly the aminoglycoside class of antibiotics, binds in the close vicinity of RafH NTD binding site (Supplementary Fig. S7). Therefore, RafH NTD would also block the binding of these antibiotics and make Mtb more drug resistant, most likely by a similar mechanism earlier observed for the aminoglycoside resistance in Mtb^31^ and in *Listeria monocytogenes*^46^.

The RafH CTD, which is longer than the HPF^long^ CTD, binds at the PBC and is sandwiched between bS1 and uS11 r-proteins. The PBC has been proposed as the binding site of mRNA before translation initiation and regulates the initiation process^47^. Therefore, the presence of RafH CTD would block its binding (Fig. 4d). Furthermore, its binding also blocks the bS1 mediated translation initiation for mRNA’s, having AU-rich sequence elements instead of AG-rich SD sequence elements in its 5’UTR regions^48^.

Previously reported structure for HPF^short^/RMF and YifA showed bS1 involvement in ribosome hibernation ^19, 27^. However, no such contribution is described for HPF^long^ mediated 100S ribosome formation^15^. Interestingly, we found that bS1 r-protein was present in a small fraction (23%) of the total particles (Supplementary Fig. S1b), suggesting its involvement in RafH mediated ribosome hibernation and further stabilizing the RafH CTD binding. Additionally, the docking studies revealed that bS1 r-protein binding site in the mycobacterial 70S ribosome (Fig. 5a) overlaps with the HPF^long^ CTD binding in 100S ribosome (Fig. 5b) thus, bS1 in mycobacteria would prevent the formation of 100S like ribosome complex, in addition to the severe steric hindrance caused by RafH CTD, H54a, and bS1 at the dimer interface (Fig. 5c).

The H54a of 23S rRNA, unique to the mycobacterial ribosome^28, 30^ adopts a different conformation and appears to interact with RafH CTD (Fig. 2b and Fig. 4a). This interaction would further strengthen the 70S stability and also suggests, a role of H54a in ribosome hibernation. A similar conformation was observed in earlier reported 70S ribosome HPF complex^32^.

The linker region, which connects the two domains, is of varying length and found to be disordered in the HPF^long^ structures reported so far^15^, with this, it was also propounded that the HPF^long^ linker might not interact with the a-SD sequence of 16S rRNA^24^. Noteworthy, we found that in mycobacteria, the RafH linker regions interact with a-SD sequence of 16S rRNA through electrostatic and base stacking interactions (Fig. 2d and Supplementary Movie 1). This binding would block the interaction between the mRNA SD sequence with the 16S rRNA a-SD sequence (Fig. 4d). This is critical for correctly positioning the mRNA in the 30S subunit during translation initiation^49^. Thus, RafH presence would also block the translation initiation of leadered (with 5’UTR) mRNA.

The presence of E-site tRNA with and without RafH in two classes (Supplementary Fig. S1) indicates that E-site tRNA binding might not influence the RafH binding. However, it may be a strategy to hoard tRNA, which would be required immediately upon restoration of active growth.

The MPY NTD, another HPF known to mycobacteria, binds to the similar RafH NTD binding site, whereas the linker region and CTD remain disordered in reported cryo-EM single particle reconstruction maps^31, 32^. Interestingly, MPY possesses amino acid residues similar to HPF^long^, which is shorter than the RafH (Supplementary Fig. S6). Likewise, HPF^long^ CTD, the two MPY CTD could dimerize and consequently induce ribosome dimerization and 100S formation. But MPY hibernates ribosomes in the 70S form only^31, 32, 37^, and its CTD structure remains unknown^31, 32^. It would be interesting to see what prevents MPY CTD dimerization once the binding site in the 70S ribosome is resolved. Perhaps, the H54a and bS1 would prevent the two ribosomes from coming close forming 100S-like ribosome dimers. However, it needs further experimental validation.

The RafH is a actinobacteria specific HPF. Thus, mycobacteria have evolved with distinctive RafH mediated ribosome hibernation exhibiting a noble way of translation inhibition, antibiotic resistance, and stabilizing the 70S structure. Therefore, the structure-based design of the modulator of mycobacterial ribosome hibernation may offer a promising strategy to prevent Mtb’s entry into the LTBI and shorten the length of TB treatment with a reduced chance of disease relapse.

## Methods

### Ribosome isolation and purification

The ribosomes from *M. smegmatis* (mc^2^155) were isolated following a similar protocol as reported earlier^50^. The cells were grown at 37°C till the mid-log phase (0.6 OD_600_) in Sauton’s media and pelleted at 8,000 rpm for 30 minutes. The cells were lysed using Mixture Mill MM500 (Retsch) for six cycles, each at 30 hertz for 1 minute in cryo-condition. A lysis buffer (20 mM HEPES pH 7.4, 20 mM MgCl_2_, 100 mM NH_4_Cl, 1 mM PMSF, 3 mM DTT, 1X Protease inhibitor cocktail) was used to resuspend the cell lysate. Cell debris was removed by centrifugation at 13,000 rpm for 30 minutes, and the clear supernatant was layered on a 1.1 M sucrose cushion in buffer A (20 mM HEPES pH 7.4, 20 mM MgCl_2_, 100 mM NH_4_Cl, 3 mM DTT) in 1:1 ratio and ultracentrifuged at 1,00,000g using rotor P70AT2 (Hitachi). The crude ribosome pellet was dissolved in buffer B (20 mM HEPES pH 7.4, 20 mM MgCl_2_, 50 mM NH_4_Cl, 3 mM DTT) and homogenized using a Dounce homogenizer followed by DNaseІ treatment (3 U/µl, ThermoFisher) for one hour on ice, subsequently, centrifuged at 13,000 rpm for 30 minutes at 4°C. The concentration of the crude ribosome in the supernatant was estimated by measuring absorbance at 260 nm. For further purification, 10-15 O.D units of crude ribosomes were layered on a 10-40% sucrose gradient in buffer C (20 mM HEPES pH 7.4, 20 mM MgCl_2_, 30 mM NH_4_Cl, 3 mM DTT). The gradients were prepared by using BioComp Gradient Master and then ultracentrifuged at 38,000 rpm for 4.5 hours (P40ST rotor Hitachi) and fractionated using a Gilson fractionator in a BioComp station (Fig. 1a). The peaks from the sucrose gradient fractionation were analyzed on 0.06% bleach agarose gel (Fig. 1a). The 30S, 50S, 70S, and polysome fractions were concentrated separately using 100 kDa Amicon (Millipore) and stored in buffer D (20 mM HEPES pH 7.4, 20 mM MgCl_2_, 30 mM NH_4_Cl, 3 mM DTT).

### Ribosome dissociation and re-association

For dissociation of the 70S ribosome to respective subunits, 30S and 50S, the MgCl_2_ concentration was reduced from 20 mM to 1 mM by passing 10 ml buffer E (20 mM HEPES pH 7.4, 1 mM MgCl_2_, 30 mM NH_4_Cl, 3 mM DTT, 0.1 mM spermidine) three time followed by incubation on ice for 3-4 hours, and finally concentrated using 100 kDa Amicon (Millipore). The ribosomes were layered on a 10-40% sucrose gradient prepared in buffer E, and ultracentrifugation was carried out at 38,000 rpm for 4.5 hours (P40ST, Hitachi). The gradients were fractionated and again analyzed on a 0.06% bleach agarose gel (Fig. 1b). The peaks corresponding to 50S and 30S were concentrated separately, and concentration was estimated by measuring absorbance at 260 nm. The 70S ribosomes were re-associated by mixing the equimolar concentration of the 50S and 30S ribosomes, and the concentration of MgCl_2_ was increased from 1 mM to 20 mM. These re-associated ribosomes were analyzed similarly by density gradient centrifugation (Fig. 1c).

### RafH overexpression and purification

The *M. smegmatis* gene MSMEG_3935 encoding RafH protein was commercially synthesized from GenScript and cloned in the pET-28a(+) bacterial expression vector with C-terminal containing His_6_-tag. The presence of MSMEG_3935 was confirmed by double digestion using Ndel and Xho1 restriction enzymes. The RafH protein was overexpressed in *E. coli* C41(DE3) cells. The cells were grown at 37°C till 0.6 OD_600_, cell culture was chilled at 4°C for 30 minutes, and then RafH overexpression was induced by adding 0.5 mM IPTG. The cells were further grown at 16°C for 16 hours at 180 rpm and pelleted at 8,000 rpm for 30 minutes. The cell pellet was stored at -80°C. The cells pellet was lysed by sonication in lysis buffer (50 mM Tris-HCl pH 7.0, 500 mM NH_4_Cl, 10 % glycerol, 20 mM Imidazole, 0.5% Tween 20, 10 mM MgCl_2_, 1 mM PMSF, Proteinase K 50 µg/ml, 5 mM β-ME). Then, the lysate was pelleted down at 13,000 rpm for 1 hour at 4°C. The supernatant was incubated with Ni-NTA (Millipore) beads for 2-3 hours on a rocking shaker at 4°C followed by 3 times washing with wash buffer (50 mM Tris-HCl pH 7.0, 500 mM NH_4_Cl, 10 % glycerol, 20 mM Imidazole, 10 mM MgCl_2_, 5 mM β-ME) to remove the non-specific bound protein. Then, protein was eluted (1ml fraction) with 20 ml elution buffer (50 mM Tris-HCl pH 7.0, 500 mM NH_4_Cl, 10 % glycerol, 300 mM Imidazole, 10 mM MgCl_2_, 5 mM β-ME). The protein fractions were pooled and concentrated in a 10 kDa cut-off Amicon filter (Millipore). Further protein purification was performed by size-exclusion chromatography using Superdex^Tm^ 200 increase 10/300 column (Cytiva). The protein purity was confirmed with SDS-PAGE (Fig. 1d). The RafH protein fraction was pooled and concentrated in a 10 kDa cut-off Amicon filter (Millipore). The protein concentration was checked by measuring absorbance at 280 nm, and protein at 1.2 mg/ml concentration was stored at -80°C.

### Ribosome RafH complex preparation and sucrose pelleting assay

The ribosome RafH complex was prepared in 100 µl reaction volume by incubating 1 µM 30S with 1 µM 50S at 37°C for 10 minutes in a complex-binding buffer (20 mM HEPES pH 7.4, 20 mM MgCl_2_, 100 mM NH_4_Cl, 3 mM DTT) followed by incubation on ice for 5 minutes.10 µM of RafH protein was added to this reaction mixture, and 10 µl of buffer was added to the control sample and incubated for 20 minutes at 37°C. 80 µl of this reaction mixture was layered on a 0.8 M (500 µl) sucrose cushion in a 1 ml open-thick wall polypropylene tube. The ribosome RafH complex was pelleted down by ultracentrifugation at 6,00,000g for 4 hours in a Beckman Coulter rotor MLA-150. The pellet was resuspended in a 50 µl complex binding buffer, and the supernatant was concentrated using 10 kDa cut-off Amicon (Millipore) till the volume reached 50 µl. Further presence of RafH was investigated by running supernatant and pellet fractions of both reaction and control samples in 12% SDS-PAGE (Fig. 1d).

### *In-vitro* translation inhibition assay

A luminescence based translation inhibition assay was performed using an *in-vitro* translation PURExpress^®^ Δ Ribosome Kit from (NEB) The constituents of the kit were incubated with 25 ng pMSR DNA template, 0.1 µl ribonuclease inhibitor (5 units), 180 nM crude ribosomes, with different molar concentrations of RafH (0 to 40X), at 37°C for 1 hour. A total 6 reactions in triplicates with 5 µl reaction volume were incubated, then the reaction was quenched by keeping the reaction mixture on ice for 10 minutes. The luminescence was measured immediately after adding 20µl NanoGlo substrate by using a GLOMAX luminometer from Agilent Technology (Promega) (Fig.1e)

### Electron microscopy

For preliminary screening, negative staining was performed. 3 µl of 1 mg/ml 70S ribosome RafH complex was applied on a glow discharged 300 CF300-Cu grids (EMS), the excess sample was blotted, washed with MilliQ water, and stained with 1% uranyl acetate solution. The grids were screened in JEOL 1400 JEM, 120 kVa microscope. The cryo-EM grids were prepared using Gatan’s CP3 plunger for cryo-EM condition optimization. 3 µl sample was applied on a glow discharged grid R 1.2/1.3 on 300 mesh Cu Quantifoil from TED PELLA, INC and blotted for 3 seconds before plunging grids into the liquid ethane. The grids were mounted on Gatan 626 Cryo-holder and analyzed in JEOL 2200 FS JEM, 200 kVa microscope equipped with the Gatan K2 Summit direct electron detector camera. The data was collected at a low dose of 1.3 e/Å^2^/frame in movie mode, 30 frames per movie stack at 1.3 Å pixel size by using JEOL’s automatic data collection software, JADAS (Fig. 1f). All the initial sample optimization and grid screening were done at the Advanced Technology Platform Centre (ATPC), Regional Centre for Biotechnology (RCB), Faridabad. The high-resolution data was collected using 300 kVa Titan Krios (Thermo Fischer) equipped with Falcon 3 direct electron detector camera at National Electron Cryo-Microscopy Facility, Bangalore Life Science Cluster (BLiSc), Bangalore. The data was collected in an electron counted movie mode. 12,343 movie stacks were collected with 25 movie frames per stack at 1.07 Å pixel with an electron dose of 1.34 e/Å^2^/frame (Supplementary Fig. S1a and Supplementary Table S1).

### Single particle reconstruction

The single particle reconstruction was carried out using Relion-3.1.4^51^. A summary of data processing is given in Supplementary Fig. S1. The movie frames were drift corrected, and single micrographs were generated using Relion-3.1.4. The micrographs were CTF corrected using CTFFIND4^52^. 12,02,461 auto-picked particles were subjected to two rounds of 2D classification, and the best 2D classes containing 7,30,969 particles were selected. These particles were subjected to 3D classification, and classes showing density for RafH, containing 3,28,619 particles, were subjected to 3D refinement. A 60 Å lowpass filtered 70S ribosome cryo-EM map (EMDB ID; 8932)^31^ was used as a reference map. A focused 3D classification on a small subunit without alignment was performed. The one class which shows apparent density for RafH CTD with 1,53,262 (47%) particles yielded a cryo-EM map at 3.0 Å resolution after 3D refinement and post-processing. The gold-standard FSC = 0.143 criterion^53^ was used for resolution estimation. The CTF refinement and particle polishing^51^ were used to further improve the resolution to 2.8 Å (Supplementary Fig. S1a).

However, the density for the RafH CTD was weaker than expected for a map at this resolution. Therefore, a focused 3D classification was performed with signal subtraction without alignment (FCwSS) with a regularization parameter of T = 12^54^. The signal for RafH CTD and its surrounding regions, H54a, bS1, and uS2, was subtracted, and FCwSS was performed by classifying into 5 classes (Supplementary Fig. S1b). Class 1, with 44,299 (28%) particles, showed fragmented cryo-EM density. Class 2 with 36,121 (23%) particles that contains RafH CTD and bS1. Class 3, with 30,514 (20%) particles, contains RafH CTD. Class 4 with 44,299 (28%) particles that contains RafH CTD and E-site tRNA. Class 5 contains 1% unaligned particles. Classes 2, 3, and 4 were subjected separately for 3D refinement and postprocessing. Final maps were interpreted by applying a 3.5 Å resolution low pass filter as RafH CTD still has a lower density than the ribosome core. The particles from these three classes, 2, 3, and 4, were joined together with a total of 1,10,934 particles, which yielded a final map of 2.8 Å resolution, upon 3D refinement and postprocessing (Supplementary Fig. S1b). The local resolution for the final maps were calculated using ResMap^55^ (Supplementary Fig. S2).

To deal with the inherent ribosomal inter subunit motion, a 3D multi-body refinement^56^ was carried out by treating LSU and SSU as two bodies with 10 Å rotation and 2-pixel translation. It has further improved the map quality of individual subunits and yielded the final resolution of 2.7 Å and 2.9 Å for LSU and SSU, respectively (Supplementary Fig. S3). Similarly, a multi-body refinement was carried out for class 2, which contains bS1, and class 4 which has E-tRNA in addition to RafH, and final maps were low pass filtered to 3.5 Å resolution (Supplementary Fig. S1b) because of poor resolution for CTD, bS1, and E-tRNA compared to the core of the ribosome.

### Model building and structure analysis

The atomic coordinates of *M. smegmatis* 70S ribosome (PDB ID; 6DZI)^31^ was docked in the final cryo-EM map using Chimera^57^. The refinement was performed using phenix.real_space_refinement^58^. The model building was carried out using COOT v.0.9.3^59^. For RafH and bS1 model building, the initial models were obtained from AlphaFold2^60^ and docked in cryo-EM map. The linker region was manually built in COOT. The final model quality was checked using MolProbity^61^. Figures were prepared in Chimera and ChimeraX^62^.

## Data availability

The atomic coordinates and the final maps are being submitted to the protein data bank (PDB) and electron microscopy data bank (EMDB).

## Acknowledgment

We acknowledge the Electron Microscopy facility at Advanced Technology Platform Centre, Regional Centre for Biotechnology, Faridabad and thank Dr. Reena and Mr. Madhava for help in initial cryo-EM sample screening. We acknowledge National Electron Cryo-Microscopy Facility, Bangalore Life Science Cluster (BLiSc), Bangalore, and special thanks to Drs. Vinothkumar Kutti and Sucharita Boss for cryo-EM data collection. We thank Dr. Todd Gray, Wodsworth Center Albany, USA, for providing the plasmid used in the *in-vitro* translation assay. N.K. & S. S. acknowledges fellowships from UGC and DBT, respectively. This work is supported by Early Carrier Research Award grant to P.S.K. from SERB DST.

## Author contribution

P.S.K. conceived this study. N. K. purified the RafH protein, S. S. purified the ribosome. N. K. prepared the ribosome RafH complex with the help of S. S. . N. K. performed the *in-vitro* translation assay. P.S.K. and N. K. collected initial data. P.S.K. and N. K. performed the single particle reconstruction. P.S.K. prepared the figures and wrote the manuscript with the inputs for N. K. and S. S..

## Notes

### Competing Interest Statement

The authors have declared no competing interest.

