## Supplementary Figures for "Cryo-EM structure of the mycobacterial 70S ribosome in complex with ribosome hibernation promotion factor RafH, reveals the unique mode of mycobacterial ribosome hibernation"

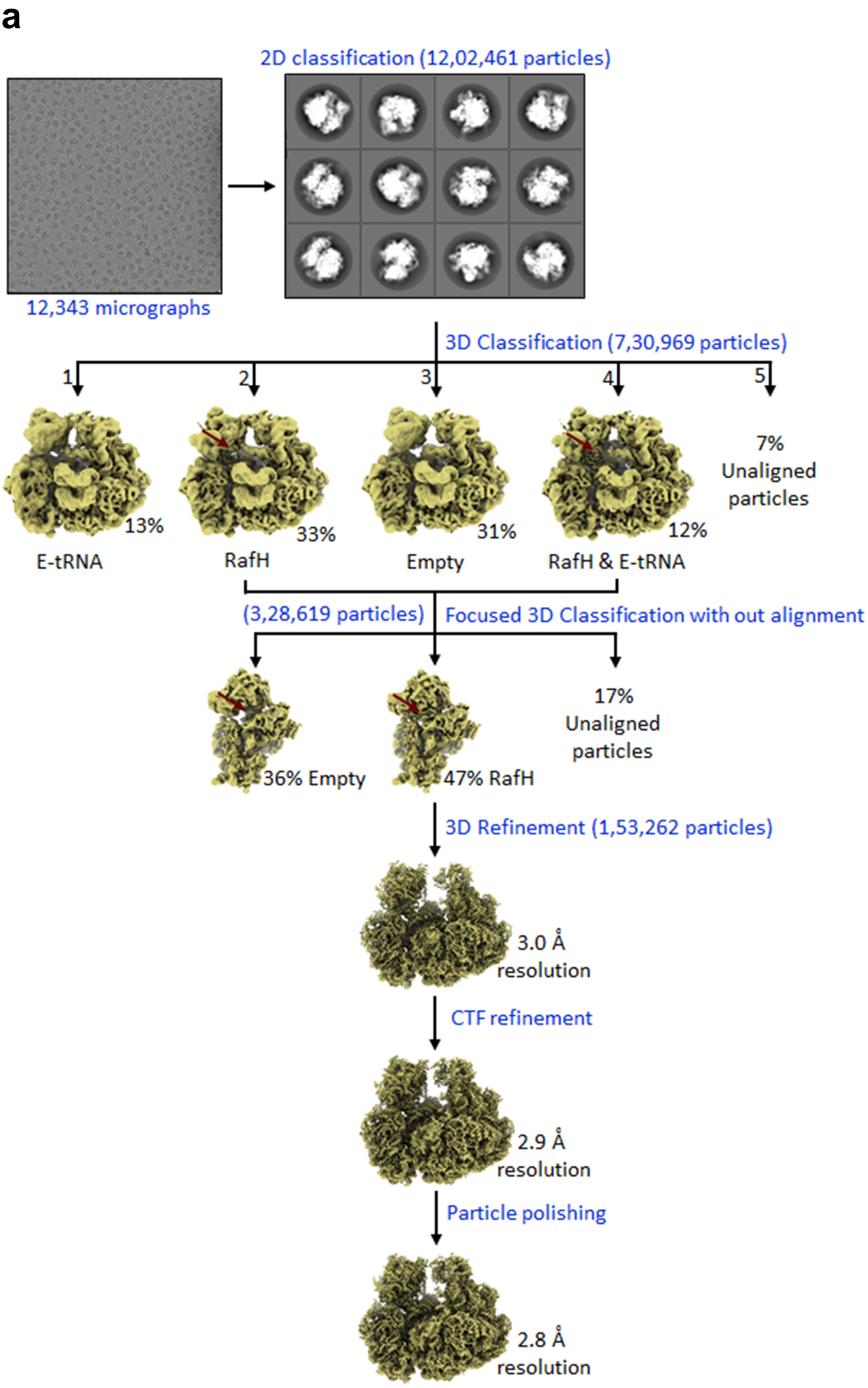

a
2D classification (12,02,461 particles)
12,343 micrographs
3D Classification (7,30,969 particles)
1
2
3
4
5
13%
33%
31%
12%
7%
Unaligned
particles
E-tRNA
RafH
Empty
RafH & E-tRNA
(3,28,619 particles)
Focused 3D Classification with out alignment
36% Empty
47% RafH
17%
Unaligned
particles
3D Refinement (1,53,262 particles)
3.0 Å
resolution
CTF refinement
2.9 Å
resolution
Particle polishing
2.8 Å
resolution

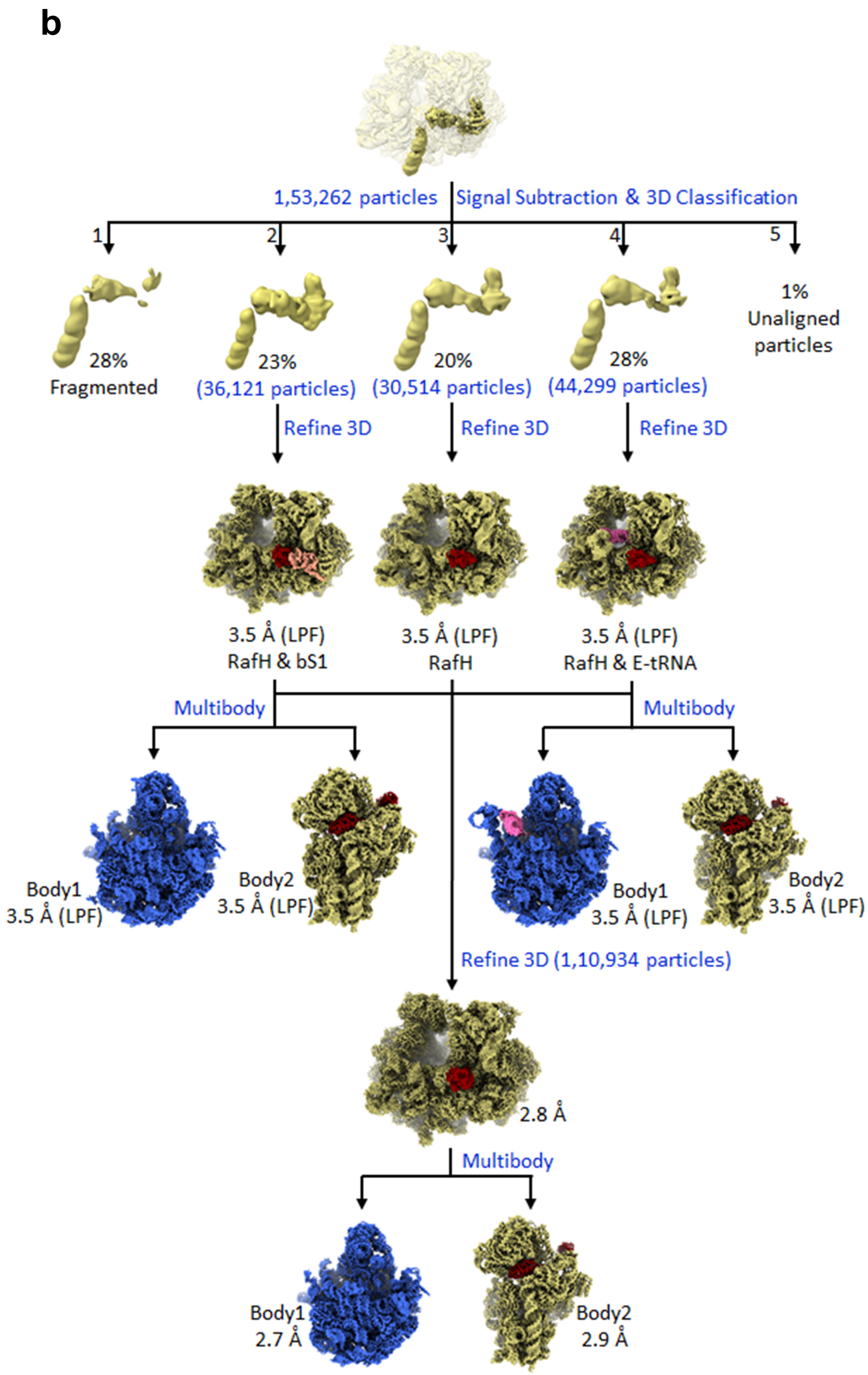
b
1,53,262 particles
Signal Subtraction & 3D Classification
1
2
3
4
5
28%
Fragmented
23%
(36,121 particles)
20%
(30,514 particles)
28%
(44,299 particles)
1%
Unaligned particles
Refine 3D
Refine 3D
Refine 3D
3.5 Å (LPF)
RafH & bS1
3.5 Å (LPF)
RafH
3.5 Å (LPF)
RafH & E-tRNA
Multibody
Multibody
Body1
3.5 Å (LPF)
Body2
3.5 Å (LPF)
Body1
3.5 Å (LPF)
Body2
3.5 Å (LPF)
Refine 3D (1,10,934 particles)
2.8 Å
Multibody
Body1
2.7 Å
Body2
2.9 Å

**Supplementary Figure 1. Summary of single particle reconstruction.** (a) 2D classification, 3D classification and initial consensus 3D maps and particle polishing are shown. The RafH NTD binding site is shown in a maroon arrow. (b) Signal subtraction with 3D classification without alignment and final multi-body refinement are shown. Maps were low pass filtered (LPF) All steps were performed using RELION 3.1.4.

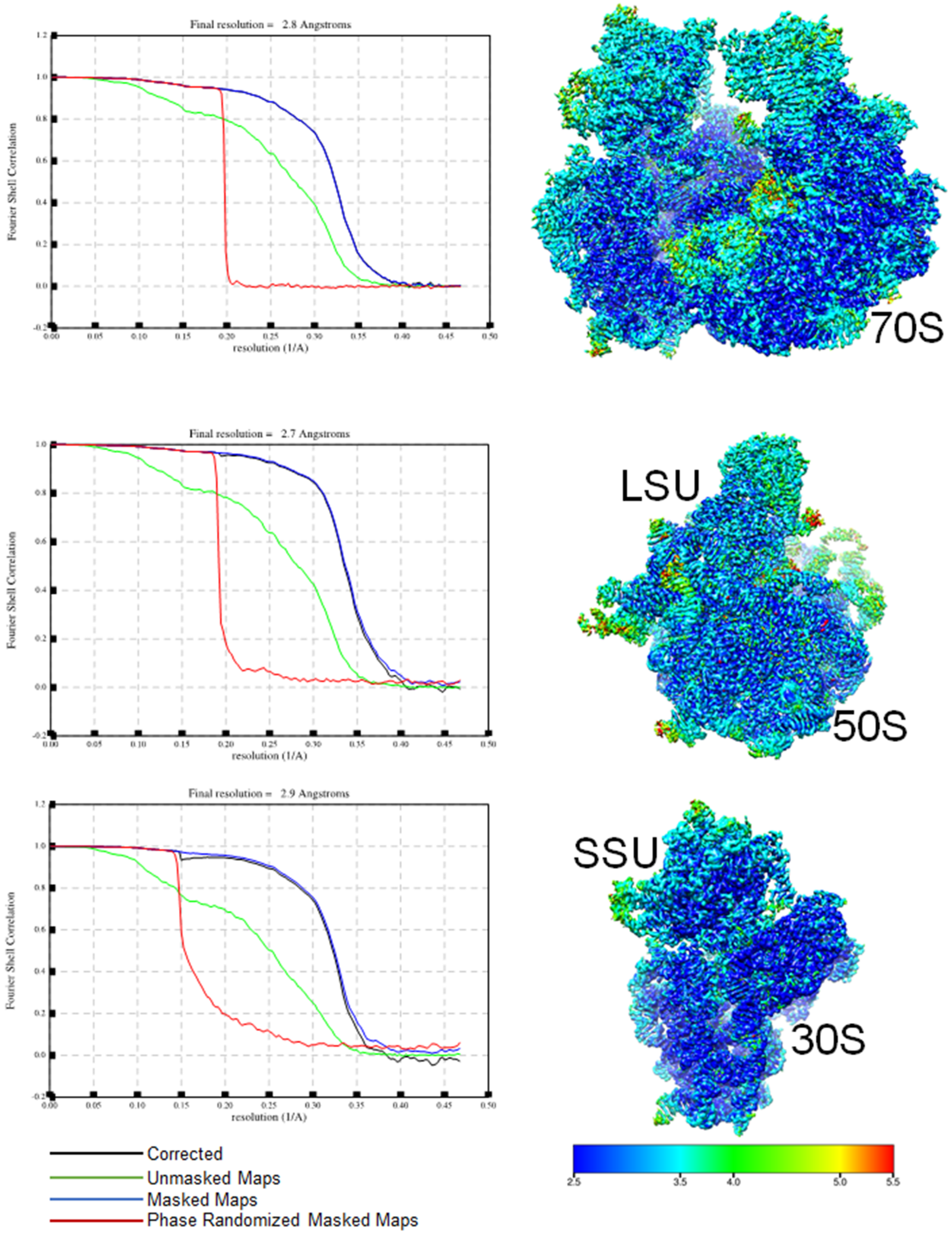


**Supplementary Figure 2 Fourier Shell Correlation and local resolution.** FSC for the 70S, 50S and 30S in top, middle, and bottom, respectively, shown in the left panel. The local resolution for the 70S, 50S, and 30S in the top, middle, and bottom, respectively, shown in the right panel. The color bar for resolution is shown in the bottom right.

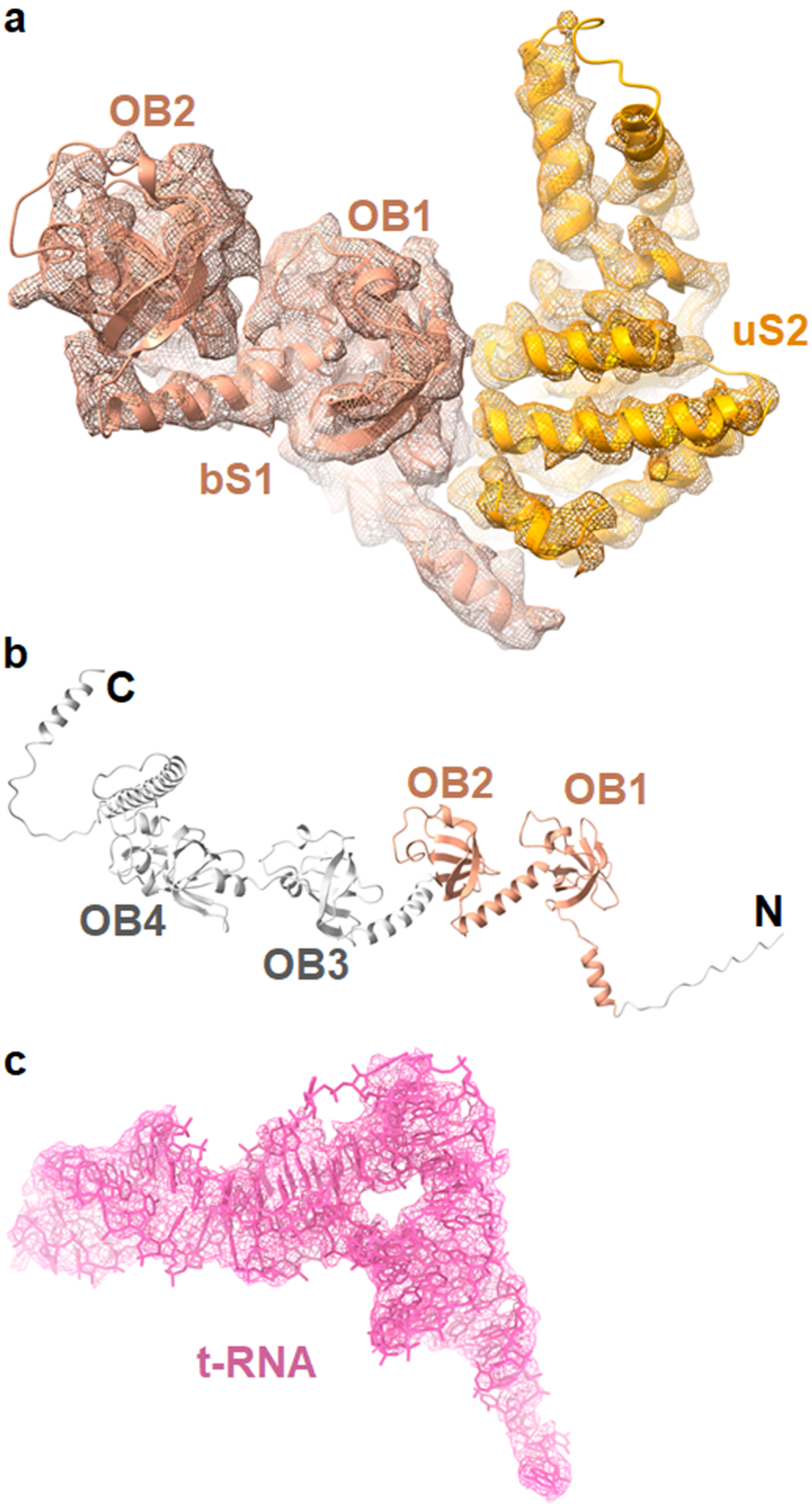


**Supplementary Figure 3 Cryo- EM density and models.** The cryo- EM density corresponds to the OB1 and OB2 domains of bS1, and uS2 in mesh, and their models in the ribbon are shown. (b) the full-length structure of bS1 predicted using AlphaFold2. The domains the OB1 and OB2 (dark salmon) and unmodeled regions (gray) are shown. (c) the E-tRNA cryo- EM density in mesh (pink) and model in sticks (pink) is shown.

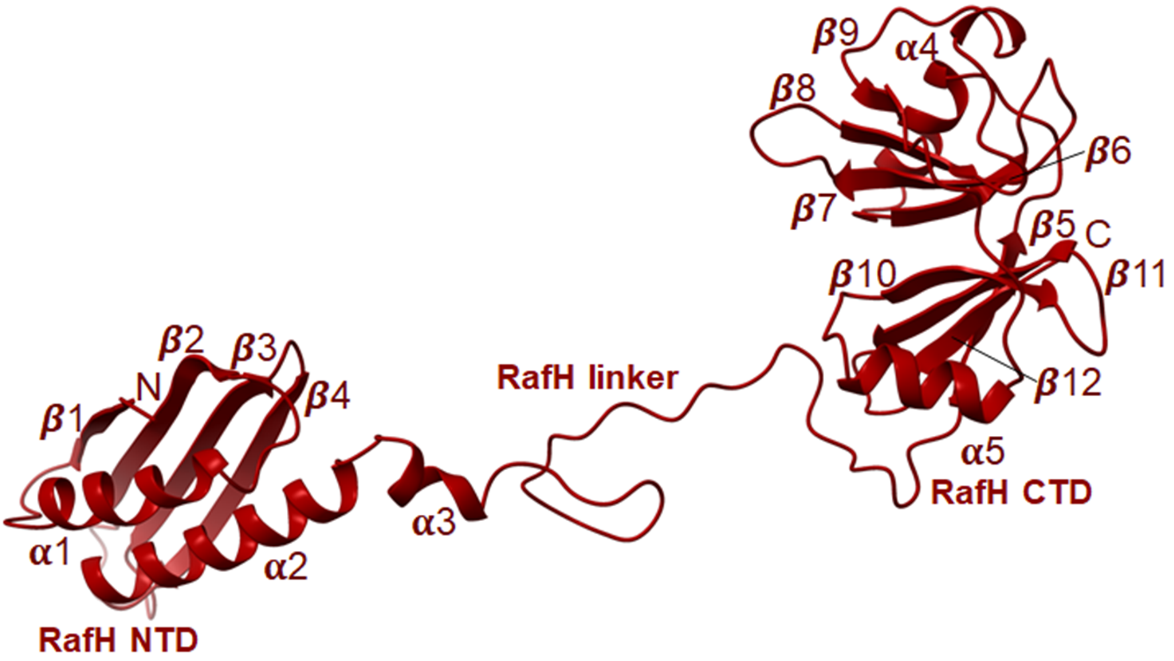


**Supplementary Figure 4 RafH full-length model.** The RafH full-length structure in ribbon style with labeled secondary structures, domains, and linker is shown.

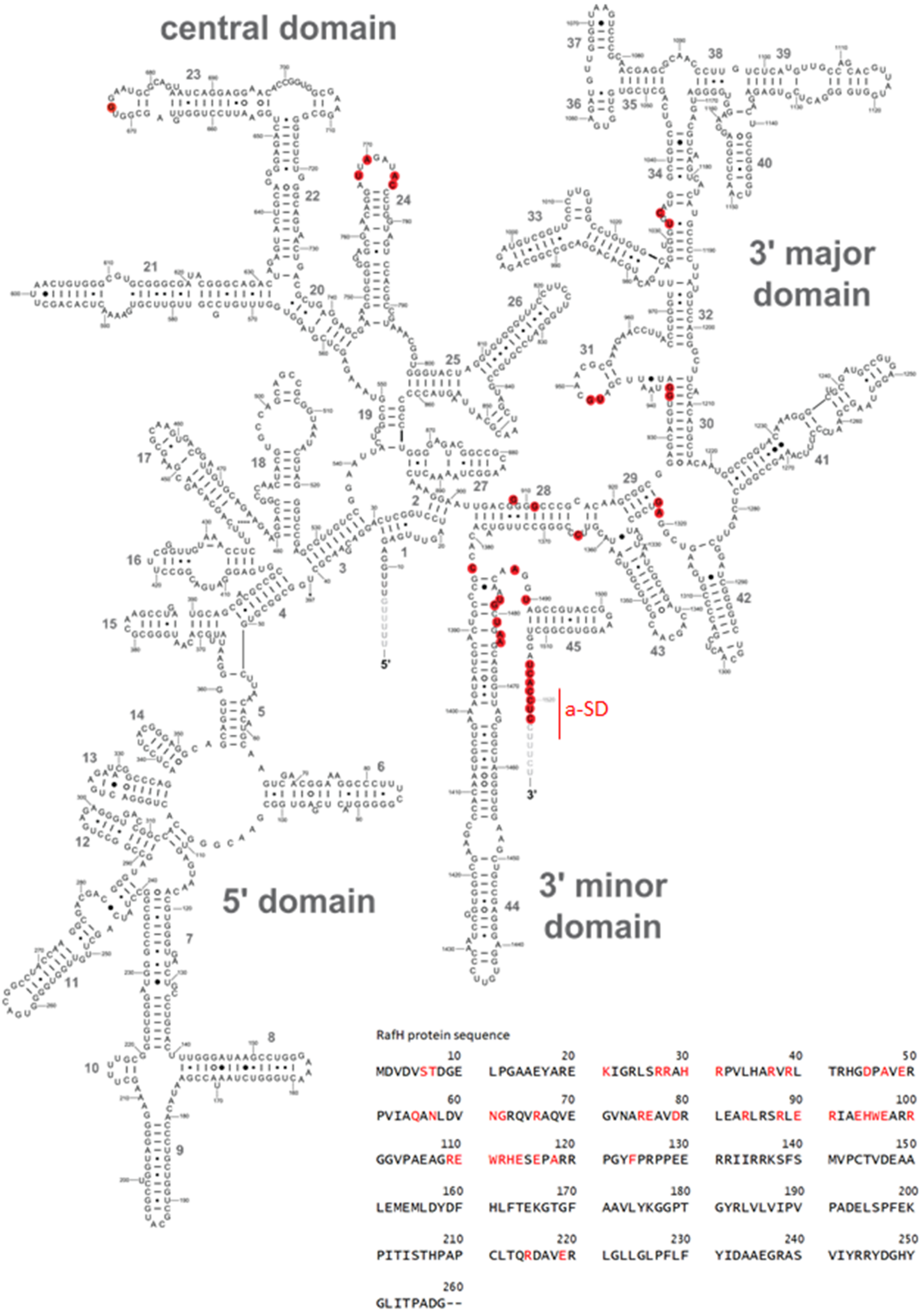


**Supplementary Figure 5 16S rRNA and RafH interaction.** The 16S rRNA 2D diagram and RafH sequence are shown. The RafH amino acid residues (red) and nucleotides (highlighted in red) are involved in interactions. The template for 16S rRNA 2D diagram was adopted from (Hentschel *et al.*, 2017).

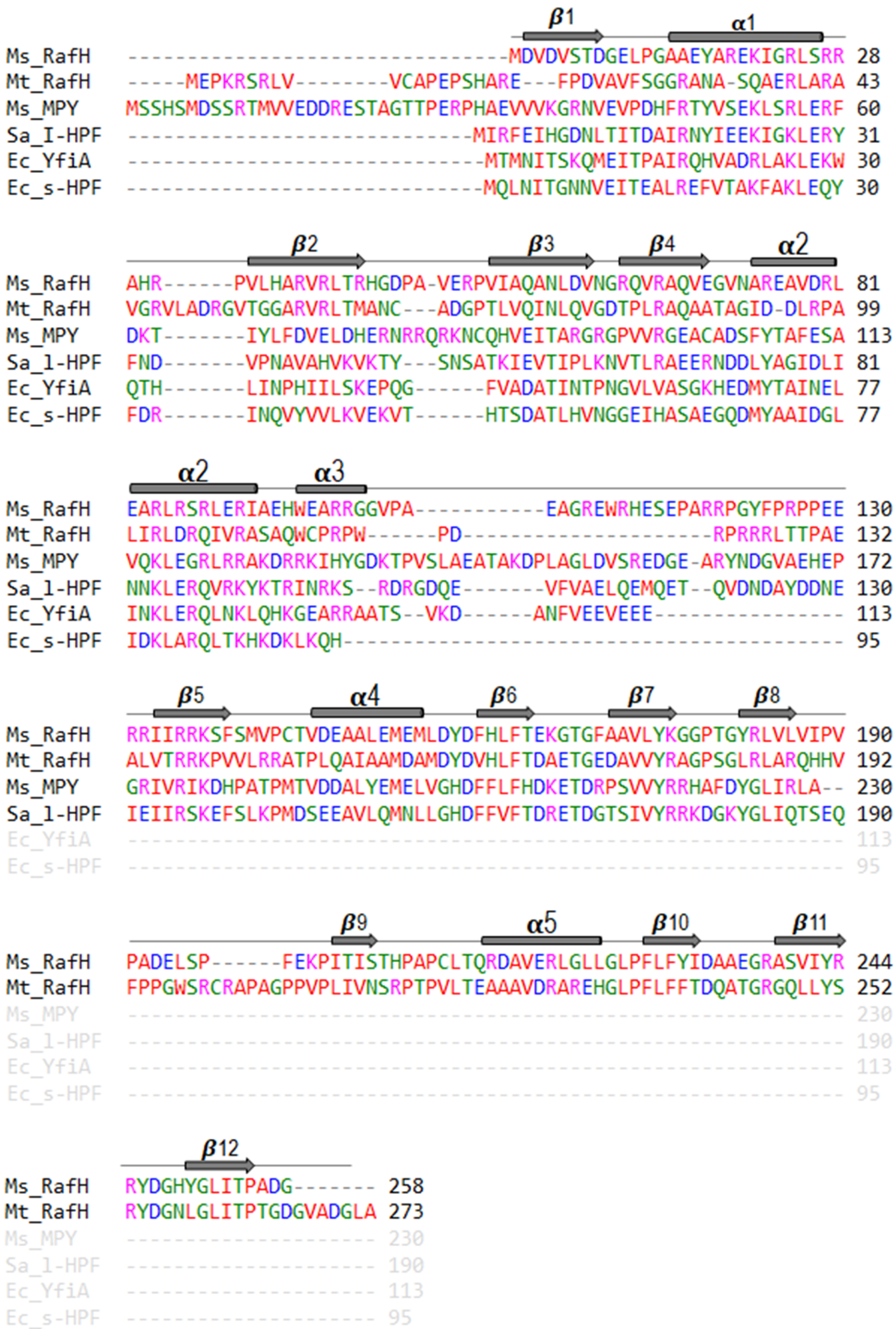


**Supplementary Figure 6 Multiple sequence alignments.** The multiple sequence alignment for *M. smegmatis* RafH (Ms_RafH), *M. tuberculosis* RafH (Mt_RafH), *M. smegmatis* MPY (Ms_MPY), *S. aureus* HPF$^{long}$ (Sa_l-HPF), *E. coli* YfiA (Ec_YfiA), *E. coli* HPF$^{short}$ (Ec_s-HPF) with the secondary structures present in RafH are shown.

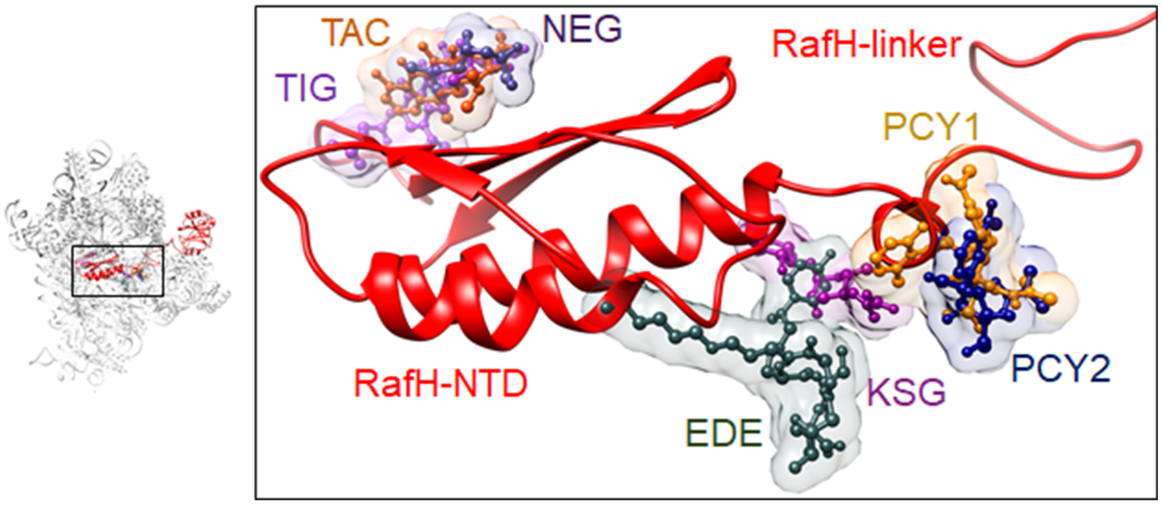


**Supplementary Figure 7 RafH NTD and antibiotic binding.** The RafH and antibiotic binding in its vicinity on 16S rRNA is shown in the thumbnail on the left side. A magnified view is shown on the right side. The antibiotics were docked onto RafH ribosome 30S. The antibiotics; Tigecycline (TIG) (PDB ID; 4V9B), tetracycline (TAC) (PDB ID; 4V9A), Negamycine (NEG) (PDB ID; 4WF1) Edeine (EDE) (PDB ID; 1I95), Pactamycin1 (PCY1) (PDB ID; 4KHP), Pactamycin2 (PCY2) (PDB ID; 4W2H) and Kasugamycin (KSG) (PDB ID; 4V4H) are shown.
